# Crossing the Blood-Brain-Barrier: A bifunctional liposome for BDNF gene delivery – A Pilot Study

**DOI:** 10.1101/2020.06.25.171264

**Authors:** Danielle M. Diniz, Silvia Franze, Judith R. Homberg

## Abstract

To achieve their therapeutic effect on the brain, molecules need to pass the blood-brain-barrier (BBB). Many pharmacological treatments of neuropathologies encounter the BBB as a barrier, hindering their effective use. Pharmaceutical nanotechnology based on optimal physicochemical features and taking advantage of naturally occurring permeability mechanisms, nanocarriers such as liposomes offer an attractive alternative to allow drug delivery across the BBB. Liposomes are spherical bilayer lipid-based nanocapsules that can load hydrophilic molecules in their inner compartment and on their outer surface can be functionally modified by peptides, antibodies and polyethyleneglycol (PEG). When composed of cationic lipids, liposomes can serve as gene delivery devices, encapsulating and protecting genetic material from degradation and promoting nonviral cell transfection. In this study, we aimed to develop a liposomal formulation to encapsulate a plasmid harbouring brain-derived neurotrophic factor (BDNF) and infuse these liposomes via the peripheral bloodstream into the brain. To this end, liposomes were tagged with PEG, transferrin, and arginine and characterized regarding their physical properties, such as particle size, zeta-potential and polydispersity index (PDI). Moreover, we selected liposomes preparations for plasmid DNA (pDNA) encapsulation and checked for loading efficiency, in vitro cell uptake, and transfection. The preliminary results from this pilot study revealed that we were able to replicate the liposomes synthesis described in literature, achieving compatible size, charge, PDI, and loading efficiency. However, we could not properly determine whether the conjugation of the surface ligands transferrin and arginine to PEG worked and whether they were attached to the surface of the liposomes. Additionally, we were not able to see transfection in SH-SY5Y cells after 24 or 48 hours of incubation with the pDNA loaded liposomes. In conclusion, we synthesized liposomes encapsulation pBDNF, however, further research will be necessary to address the complete physicochemical characterization of the liposomes. Furthermore, preclinical studies will be helpful to verify transfection efficiency, cytotoxicity, and in the future, safe delivery of BDNF through the BBB.

## Introduction

The development of new treatments for curing neurological diseases has been slow due to the inability of large molecule pharmaceuticals, such as products of biotechnology, recombinant proteins or gene therapies, to cross the blood-brain barrier (BBB) (Pardridge, 2020). The central nervous system (CNS) presents a series of barriers to protect itself from invading pathogens, neurotoxic molecules, and circulating blood cells. These structures with diverse degrees of permeability include the blood-brain barrier, which is the most extensive and exclusive barrier (Saraiva et al., 2016). The blood-brain-barrier (BBB) is an anatomical and physiological barrier that is responsible for the tight regulation of the transportation of cells, molecules, and ions between the periphery and the brain. Because the selective molecular permeability of the BBB prevents most of the drugs from entering into the brain, the development of new treatments for brain diseases is complicated (Pardridge, 2012). The highly specialized endothelial cells layer has an intimate contact with brain cells, so that to reach those cells, substances such as drugs, have to present suitable lipophilicity, size, and capacity to evade active extrusion (Serlin et al., 2015). The occurrence of specific receptors on the surface of the BBB facilitates the transport of various essential molecules into the brain. Nanotechnology offers an exciting approach for improving the therapeutic management of CNS diseases. Liposomes, being functionally versatile, can be engineered for targeting these receptors, thus rendering them as promising carriers for drug and gene delivery (Sharma et al., 2012). A liposome is a kind of non-viral vector that can deliver molecules such as DNA, proteins, and drugs to the action sites, and it has been used to carry exogenous gene productions (Ganly et al., 2013). They have the great advantage of being made from a natural biodegradable lipid bilayer, which is similar to the animal cell membrane structure. Thus, considering that the morphologic appearance of liposomes resembles the natural cell membrane, they are an ideal drug-carrier system (Bozzuto, 2015).

Bangham and Horne were the pioneers in reporting the synthesis of liposomes, described by them as single or multiple concentrically organized lipid bilayers containing an inner aqueous compartment (Bangham and Horne, 1964) (Figure 1). Liposomes usually contain phospholipids as its basic constitution. This lipid molecule is made up of a hydrophilic head that interacts with aqueous solutions and two hydrophobic fatty acids chains that have an affinity with each other. In aqueous solution, due to these amphiphilic characteristics, a lipid bilayer is formed creating a lipophilic inner compartment that acts as a permeability barrier, both inward and outward (Bozzuto, 2015). Liposomes can serve as gene carriers, being then called lipoplexes. Lipoplexes have enormous potential to deliver plasmids into target cells; they contain cationic lipids that due to its positive charge, can complex with negatively charged DNA molecules. Theoretically, the electrostatic interaction promotes neutralization and enhances cell-membrane-DNA communication and transfection efficiency (Parker et al., 2003). However, the positive charges on the surface can lead to nonspecific interaction with plasma and other extracellular proteins, decreasing the transfection efficiency (Urtti et al., 2000). Thus, to increase transfection, neutral or helper lipids, such as dioleoylphosphatidylcholine (DOPE), are used allow endosomal escape and plasmid dissociation from liposomes before degradation in lysosomes. (Parker et al., 2003).

**Figure 1.**
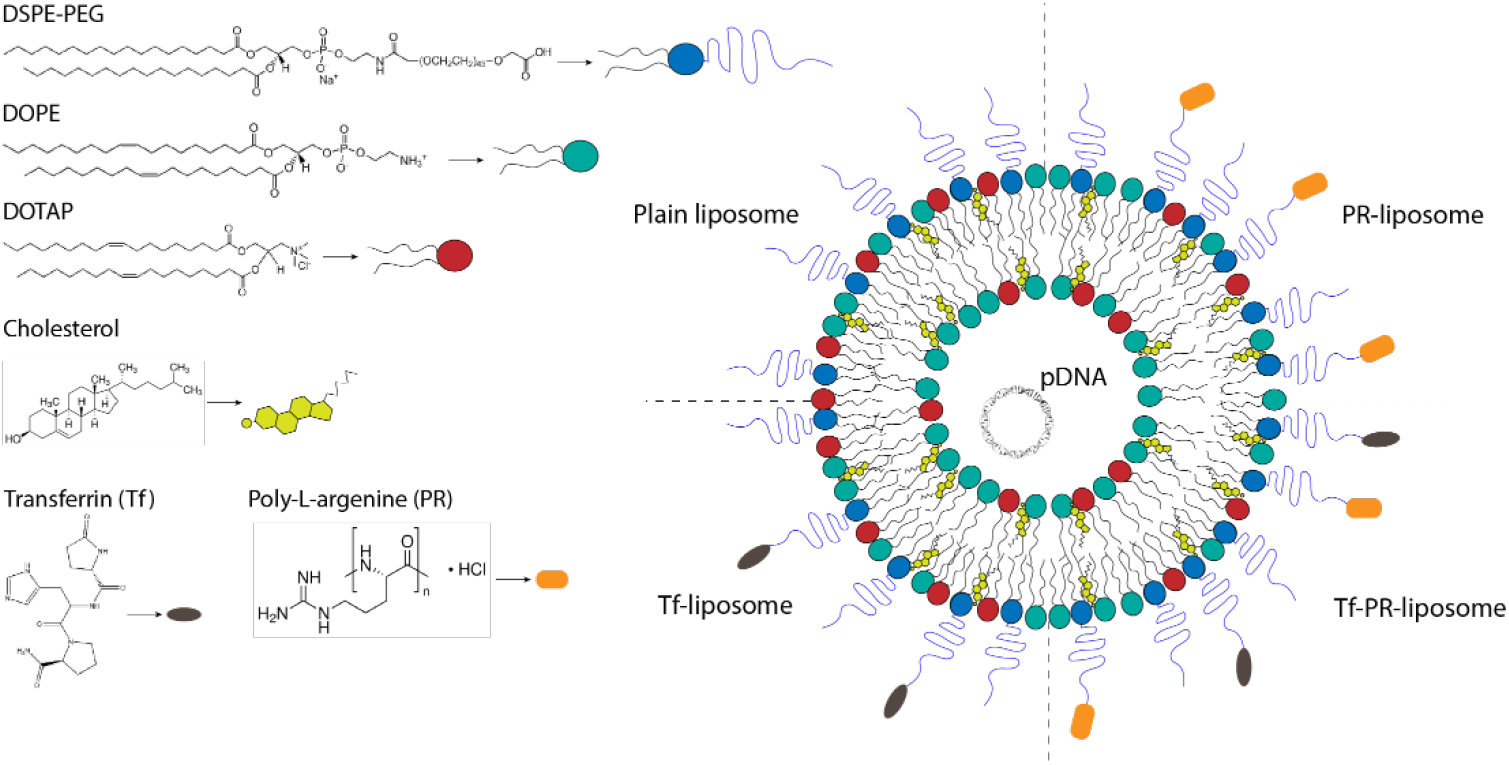
Schematic representation of the bifunctional liposomes. The structure of the lipids composing the liposome bilayer, the theoretical organization of the functional targeting with transferrin and/or arginine, and the encapsulation of the plasmid DNA (pDNA) are considered. According to Sharma et al. (2012), the following formulations were developed: Plain liposome (DOTAP/DOPE/DSPE-PEG/Cholesterol 45:45:8:2 mol %), PR-liposome (DOTAP/DOPE/DSPE-PEG/PR–PEG /Cholesterol 45:45:4:4:2 mol %), Tf-liposome (DOTAP/DOPE/DSPE-PEG/Tf–PEG /Cholesterol 45:45:4:4:2 mol %), Tf-PR-liposomes (DOTAP/DOPE/PR-PEG/Tf–PEG/Cholesterol 45:45:4:4:2 mol%).

To specifically target brain delivery, liposomes can undergo surface modifications to increase transfection efficiency. For instance, adding polyethylene glycol (PEG), or PEGylating the liposome, increases transfection efficiency, reduces toxicity, and imparts “stealth” properties (Rip et al., 2014). A widely explored liposome surface modification for brain delivery relies on the use of ligands of receptors on the brain endothelium. The cell-penetrating peptides (CPPs) are short cationic or amphipathic peptides that have the ability to transport the associated molecular cargo (e.g., peptides, proteins, oligonucleotides, liposomes, nanoparticles, bacteriophages, etc.) inside the cells (Sharma et al., 2012). Arginine, a CPP, can enhance the cellular uptake and delivery across the BBB (Morris and Labhasetwar, 2015). Moreover, the use of transferrin (Tf) is a classic method for enhancing BBB crossing via receptor-mediated endocytosis (Chen et al., 2016; dos Santos Rodrigues et al., 2020; Sharma et al., 2013, 2012; Sonali et al., 2016). In a recent study, dos Santos Rodrigues et al. (2020) successfully designed a dual-modified liposome containing CPP and transferrin ligands that promoted efficient delivery of a plasmid DNA in vitro and, into the mice brain.

Taking into consideration the large treatment inefficacy burdening individuals suffering from neuropsychiatric diseases, such as is the case of depression (Fornaro et al., 2014), the use of nanotechnology can widen up the possibilities for novel treatments. One attractive candidate is the brain-derived neurotrophic factor (BDNF), which is widely expressed in the CNS, especially in the hippocampus (Lessmann et al., 2003; Leβmann and Brigadski, 2009), and plays an important role in the regulation of several biological processes, including neuronal survival, differentiation, growth and plasticity (Alsina et al., 2001; Benarroch, 2015; Edelmann et al., 2014; Poo, 2001). It has been shown that BDNF is decreased in depressive patients (Dwivedi et al., 2003) and that antidepressant treatment increases BDNF levels (Fernandes et al., 2015; Polyakova et al., 2015; Sen et al., 2008). Local lentivirus BDNF infusion in the hippocampus alleviated depression-like behaviors in a rat model of post-stroke depression (Chen et al., 2015). Our studies in the SERT^-/-^ rat model, which presents both decreased BDNF levels in the HIP and depression- and anxiety-like behaviors (Calabrese et al., 2015; Guidotti et al., 2012; Molteni et al., 2010), showed that BDNF lentivirus infusion meliorated their anxiety levels in the novelty-induced locomotor activity. Moreover, BDNF overexpression in the hippocampus of the SERT^-/-^ rats, increased sucrose preference and intake in the sucrose consumption test, showing effects in anhedonia (Diniz et al. unpublished data). Despite promising results, BDNF local infusion into the human brain to treat depression seems to be not feasible. Moreover, neurotrophins, including BDNF do not readily cross the BBB (Pardridge, 2015). Therefore, the development of a delivery system to increase BDNF in the brain following a less invasive administration route might provide the opportunity to explore the enormous therapeutic potential of BDNF.

The present study sought to assess the feasibility to design a cell penetrating peptide tethered bi-ligand liposome for brain delivery of BDNF plasmid (Kim et al., 2014; Sharma et al., 2013, 2012). In this pilot experiment liposomes without any tag (plain liposomes), liposomes tagged with transferrin (Tf-liposomes), tagged with arginine (PR-liposomes), and bifunctional liposomes (Tf-PR-liposomes) were developed and characterized for the main physical properties, such as particle size, zeta-potential, polydispersity index (PDI) was done (see schematic representation in figure 1). Moreover, selected liposomes preparations were used for pDNA encapsulation and checked for loading efficiency, in vitro cell uptake and transfection. Pilot synthesis and characterization of the liposomal delivery system aimed to assess in first instance the efficiency of BDNF plasmid encapsulation, pursuing to the further development of a non-viral system to mediate delivery across the BBB following peripheral blood stream infusion route. mRNA BDNF overexpression in the CNS is a promising approach to remediate the decreased BDNF protein levels encountered in neurological diseases such as mood disorders.

## Material and Methods

### Materials

1,2-distearoyl-sn-glycero-3-phosphoethanolamine-N-[carboxy(polyethylene glycol) 2000] (DSPE-PEG_2000_-COOH), 1,2-Dioleoyl-sn-glycero-3-phosphoethanolamine (DOPE), and 1,2 Dioleoyl-3-trimethylammonium-propane chloride (DOTAP) were purchased from Avanti Polar Lipids (Alabaster, Alabama). Poly-L-arginine hydrochloride (molecular weight = 13,300 Da), holo-Transferrin human (Transferrin), 3β-Hydroxy-5-cholestene (Cholesterol), N-(3-Dimethylaminopropyl)-N’-ethylcarbodiimide hydrochloride (EDC), and N-Hydroxysuccinimide (NHS) were procured from Sigma–Aldrich Company (Darmstadt, Germany). The fluorescent dye 1,1’-dioctadecyl-3,3,3’,3’-tetramethyl-indocarbocyanine iodide (DiR) was obtained from Invitrogen (Landsmeer, Netherlands). BDNF (NM_012513) Rat Untagged plasmid was purchased from Origene (OriGene Technologies GmbH, Germany). All other chemicals used in this study were of analytical reagent grade.

### Methods

#### Coupling poly-L-arginine (PR) to DSPE-PEG

To obtain PR-liposomes, first, poly-L-arginine (PR) had to be coupled to DSPE-PEG. The synthesis procedure of PR–PEG was based on the previously described method (Huang et al., 2002). Briefly, PR was dissolved in 25 ml of 50mM sodium tetraborate buffer (STBB, pH 8.5) per gram of PR. The resulting solution was stirred vigorously for approximately 30 min and subsequently filtered through a 0.22μm Durapore^®^ membrane (Sterile Millex GV, Sigma–Aldrich, Buchs, Switzerland) into a sterile culture tube. The appropriate stoichiometric amount of DSPE-PEG powder was then slowly added to the solution while it was continuously stirred. After another 6h of vigorous stirring at room temperature, the solution was transferred to a dialysis tube (Spectr/Por dialysis tubing, M.W.C.O. of 6–8 kDa, Spectrum Laboratories, Inc., Rancho Dominguez, CA). The synthesized product was dialyzed out for 24h in 10mM phosphate-buffered saline (PBS, pH 7.0), followed by an additional 24 h of dialysis in deionized water. The product was then freeze-dried for 48h at −70 °C with a pressure of 0.2 mbar.

#### Verification of PR-PEG coupling

Coupling efficiency was analyzed through size-exclusion high-performance liquid chromatography performed using Agilent 1100 series HPLC system (Agilent Technologies Inc., California, USA). Two tandem Shodex Protein KW803 (8.0mm×300mm, 5μm) and KW804 (8.0mm×300mm, 7μm) columns were used at ambient temperature. The mobile phase is an aqueous solution containing 20mMHEPES buffer at pH 6.5 (prepared by dissolving 4.42 g of HEPES and 0.38 g of HEPES sodium salt in 1000mL of water and filtered through a 0.45μm filter before use), and the flow rate was 0.5 mL/min. Stock solutions of 1mg/mL PR-PEG were injected. The HPLC trace was monitored with a UV/vis detector at a wavelength of 214nm and with a RI detector at an attenuation of 7.8 × 10^3^.

#### Preparation of plain and PR-liposomes

Liposomes were prepared by lipid film hydration method as previously reported (Kim et al., 2010; Sharma et al., 2012). In short, stock solutions of each lipid in an organic solvent mixture (CHCl_3_:MeOH= 2:1, v/v) were mixed in the following molar ratio: DOTAP/DOPE/DSPE-PEG/Cholesterol 45:45:8:2 and DOTAP/DOPE/DSPE-PEG/PR–PEG/Cholesterol 45:45:4:4:2 mol/mol % for plain and PR-liposomes, respectively. The organic mixture was removed on a rotary evaporator under reduced pressure with the temperature of water bath adjusted to 40 °C. Lipid film was hydrated with PBS buffer (pH 7.4) by vigorous vortexing. The resulting liposome dispersion was extruded 10 times (Avanti^®^ Mini-Extruder, Avanti Polar Lipids, Inc.) using 0.2 μm polycarbonate membranes.

#### Development of Tf-PR liposomes

Bifunctional liposomes were prepared using the post-insertion technique (Sharma et al., 2012; Visser et al., 2005). Transferrin (Tf) was coupled to the phospholipid DSPE–PEG–COOH as reported previously (Li et al., 2009). Briefly, DSPE–PEG–COOH (remaining 4mol % of the total phospholipid content) was suspended in HEPES-buffered saline (pH 5.0) to form micelles. The micellar suspension was then treated with 360 μL of both EDC (0.5 M in H_2_O) and NHS (0.5 M in H_2_O) per 10 μmol of the phospholipid. Excess EDC was removed by dialysis, and the pH of the micellar suspension was adjusted to 7.3 with 0.1 N sodium hydroxide. Tf (125 μg/μmol of the lipid) was added to the resulting suspension and stirred at 25°C for about 8h. The resulting Tf micelles were stirred overnight with PR liposomes at room temperature, and the final liposomal dispersion was passed through a Sepharose^®^ CL-4B column to remove unbound protein.

#### Protein assay and Tf-binding efficacy

The average amount of transferrin conjugated to the liposome was quantified as described by (Anabousi et al., 2005). Shortly, one hundred microlitres of liposome dispersion were added to 400 μl of methanol. The mixture was vortexed and centrifuged (10s at 9000 x g). Then, 200 μl of chloroform were added, followed by vortexing and centrifugation (10s at 9000 x g). For phase separation, 300 μl of water were added, and the sample was vortexed again and centrifuged for 1 min at 9000 x g. The upper phase was carefully removed and discarded. Three hundred microliters of methanol were added to the chloroform phase and the interphase with the precipitated protein. The sample was mixed and centrifuged to pellet the protein (2 min at 9000 x g). Then, the supernatant was removed, and the protein pellet was dried under a stream of air. The pellet was then dissolved in 20 μl of PBS (pH 7.4), and the concentration was determined with a bicinchoninic acid (BCA) protein assay using pure holo-transferrin as standard. The coupling efficiency was calculated as mg Tf/mmol PL.

#### BDNF plasmid transformation and purification

In order to have enough pBDNF for the liposomal formulations, the pBDNF amount was increased through bacterial transformation into E. coli (DH10β), which was performed using the heat shock method (Froger and Hall, 2007). In short, after a 10 minutes incubation in ice, the mixture of bacteria and pDNA was placed at 42°C for 90 seconds (heat shock) and then placed back in ice. LB media was added, and the transformed cells were incubated at 37°C for 45-60 min with agitation. To check that isolating colonies are irrespective of transformation efficiency, two quantities of transformed bacteria were plated. The number of pBDNF copies were quantified by OD_600_ using an UV-visible spectrophotometer (NanoDrop™, USA).

After achieving the desired amplification, pBDNF was purified using the endotoxin-free plasmid purification NucleoBond^®^Xtra system (Macherey-Nagel, Germany). The final concentration was measured by OD_260_ using an UV-visible spectrophotometer (NanoDrop™, USA), and diagnostic restriction digestion was used to confirm the rough structure of the plasmid on agarose gel electrophoresis 1% TEB buffer.

#### pDNA encapsulation

Plasmid DNA was incorporated at N/P ratio of 5 into the formulation using the encapsulation method or the complexation method as previously described (Sharma et al., 2012). Shortly, in the encapsulation method, plasmid DNA was added to the hydration buffer, which was added to the lipid film and vigorously vortexed at room temperature for about 20 minutes, followed by extrusion and size exclusion column (SEC) purification (Sharma et al., 2012). In the complexation method, pDNA was added after liposomes were extruded, gently mixed and incubated for about 30 minutes to allow the lipoplex formation. Thereafter complexed liposomes were purified through SEC (Kim et al., 2010).

### Characterization of liposomes

#### Particle size and zeta potential measurements

Dynamic light scattering (DLS) is a technique in physics that can be used to determine the size distribution profile of small particles in suspension or polymers in solution (Pecora, 2000). DLS measures the time-dependent fluctuations in the intensity of scattered light, which occurs because particles (liposomes) in a suspension undergo random Brownian motion due to collisions between suspended particles and solvent molecules (Bozzuto, 2015). Therefore, the DLS allowed us to analyze the particle size and polydispersity index (PDI).

The ζ-potential is the overall charge of a particle acquires in a particular medium. A Laser Doppler Micro-electrophoresis was used to measure zeta potential. An electric field is applied to a solution of molecules or a dispersion of particles, which then move with a velocity related to their zeta potential. This velocity is measured using a patented laser interferometric technique called M3-PALS (Phase analysis Light Scattering), enabling the calculation of electrophoretic mobility, and from this, the zeta potential and zeta-potential distribution.

The size distribution, the average hydrodynamic particle size, and the zeta potential of the liposomes were evaluated using both systems described above, which are incorporated in the Zetasizer Nano (Malvern Instruments, UK). The samples were diluted in PBS-buffered saline (pH 7.4), and transferred into a disposable cuvette for particle size analysis and a capillary cell for ζ-potential measurement and inserted into the Zetasizer (Nano-ZS, Malvern Instrument, UK) at 25°C.

#### Measurement of plasmid BDNF Encapsulation Efficiency

The pBDNF encapsulation efficiency of the liposomes was calculated based on the previously reported method (Fillion et al., 2001; Gonçalves et al., 2004). DNA content of the samples was analyzed through a spectrophotometer. Shortly, DNA concentration was estimated by measuring the absorbance at 260nm, adjusting the A_260_ measurement for turbidity (measured by absorbance at 320nm), multiplying by the dilution factor, and using the relationship that an A_260_ of 10 = 50 μg/mL pure dsDNA.

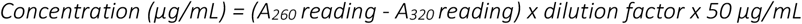

#### Cell cultures

SH-SY5Y cells were cultured in Dulbecco’s modified Eagle’s Medium (DMEM, Life Technologies), with 7.5% Fetal calf serum (FCS), 400U/ml penicillin and 100μg/ml streptomycin (Gibco), at 37°C in a 5% CO_2_ incubator. The cells were passed once a week, in a 1:5 dilution.

#### Evaluation of In-vitro Transfection Efficiency

Cells were cultured for 24 hours in a 12 wells plate, 10^3^ cells/well. After these 24 hours, 1000nM of liposomes containing GFP plasmid was added. The medium was refreshed after 1 hour. The cells were kept in an Evos FL Auto 2 cell imaging system (Invitrogen). Hourly images were automatically captured to visualize the GFP expression. The cells were kept inside the Evos at 37°C in 5% CO_2_.

## Results and discussion

### Resolving free DSPE-PEG from PEG-PR conjugate

The synthesis of the conjugate PEG-PR was particularly challenging. Because of the anticipated difficulty in revolving free PEG and PEG-conjugate, our initial efforts were focused on the identification of conditions separating these two species. We had to use different kinds of dialysis membranes to purify not only from free PEG, but also from free poly-L-arginine. During the freeze-drying procedure, we observed that the PEG-PR sample did not present loss of weight, indicating that the water molecules were not removed from the sample, and resulting in precipitation/ flocculation when we attempted to dilute PEG-PR in an organic solvent for the liposome preparation. According to Radaev and Sun (2002), during freezing, PEG tends to crystallize. This could explain increased water retention by the system (Tattini et al., 2005), and therefore solubility problems. Following the protocol proposed by Kim et al. (2010), we freeze-dried the PEG-PR solution for 48h at −70 °C with a pressure of 0.2 mbar. We also tried to decrease the temperature to −86°C and the pressure to 0.006 mbar; however, we encountered the same solubility problem.

Moreover, when measuring PEG-PR coupling efficiency through HPLC, we injected samples of 1mg/mL solutions of PR, PEG, and PEG-PR. We were not able to see a clear peak of the PR, PEG, or the PR-PEG conjugate. Altogether, we conclude that further studies are necessary to characterize the PEG-PR conjugation better. For example, through differential scanning calorimetry (DSC), the effects of the freeze-drying method over the structural and phasic changes in the conjugation of PEG with PR can be determined (Tattini et al., 2005). Additionally, Kim et al. (2010) confirmed the synthesis of PEG-PR through ^1^H NMR in D_2_O and gel permeation chromatography (GPC). Therefore, we will have to repeat the experiments and perhaps the use of other techniques could help in the identification of the coupling efficiency of the PEG-PR. The samples produced that achieved organic solvent solubility were used to develop the liposome formulations.

### Tf-binding efficacy

We planned to access the amounts of transferrin using a bicinchoninic acid (BCA) assay following Anabousi et al. (2005). We did not conduct the entire experiment, but following the specifications of the manufacturer (BCA™, Scientific, USA), we built a calibration curve using human transferrin, which would be used as the standard to analyze the transferrin content in the liposomes. The BCA assay offers the advantage of producing a linear response curve. This response curve allows accurate determination of unknown protein concentrations in the liposome surface. Given that the PEG-Tf micelles contained transferrin concentration of about 58μg/mL, which are post-inserted to the already formed and extruded liposomes, and giving the possibility that not all chains of PEG are incorporated to the final Tf-liposome, we built the curve with concentrations starting from 5μg/mL. The linear response curve (R^2^ > 0.95) obtained is showed in **figure 2** and it will help us to further determine the amount of transferrin in the liposomes.

**Figure 2.**
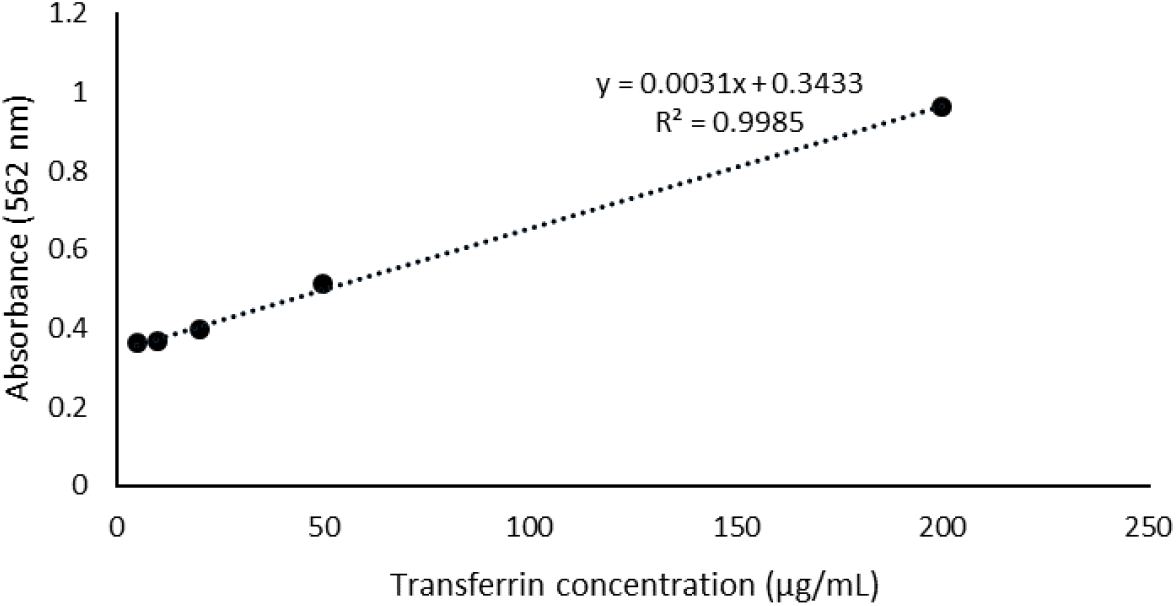
Standard curve for transferrin quantitation. Holo transferrin human in 0.9% saline (5–200 μg/mL) was used to generate standard curves for the BCA Protein Assay. The assay was conducted according to the manufacturer’s protocols in a test-tube format.

### pBDNF purification

Plasmid BDNF purification was checked by UV spectroscopy using the ratio of A_260_/A280 and A_260_/A_230_. We verified a ratio of A_260_/A280 = 1.84 ± 0.01 and A_260_/A_230_ = 2.3, indicating that the obtained samples contained a pure plasmid DNA (NucleoBond^®^Xtra, Macherey-Nagel, Germany). **Figure 3** demonstrates the result of the diagnostic restriction digestion. According to the manufacturer, the pCMV6-Entry plasmid backbone plus the BDNF insert (NM_012513) has about 5.6 kb (OriGene Technologies GmbH, Germany). We notice that digestion with the restriction enzyme Hind III should have linearized both the supercoiled and nicked forms of the plasmid, but there has probably been an incomplete digestion. Therefore, although spectroscopy analysis indicates that we obtained copies of the pBDNF, to check the size and the characteristics of the plasmid correctly, we will probably need to increase the enzyme and/or increase the incubation time to ensure complete digestion.

**Figure 3.**
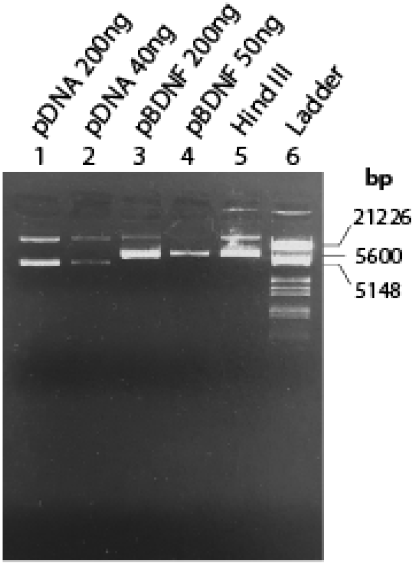
Evaluation of the pBDNF amplification and purification in agarose gel 1% TEB. Standard plasmids of known concentration were used (lanes 1 and 2), and two different concentrations were used to check uncut pBNDF (lanes 3 and 4). Digestion of 250ng of pBDNF by Hind III is expressed in lane 5, and marker III (Roche) was used as DNA molecular weight marker.

### Particle size, zeta potential and PDI characterization of the liposomes

Determining the physicochemical properties of the liposomes is essential because it can assess its passage mechanism across the BBB (Saraiva et al., 2016). There is evidence for a inverse correlation between BBB penetration and the size of the liposome (Etame et al., 2011; Hanada et al., 2014; Sonavane et al., 2008). The preferred size for brain delivery is 100nm or smaller, but studies have shown that liposomes from 100 to 140 nm have certain advantages, such as a longer half-life in blood circulation and avoidance of plasma proteins (Ross et al., 2018). Moreover, because the BBB is negatively charged, cationic liposomes can trigger cell internalization through electrostatic interactions (Harilal et al., 2020). The physico-chemical characterization of the main properties of prepared liposomes are reported in table 1.

**Table 1.** Particle Size, polydispersity index (PDI), and zeta (ζ)-potential of liposomes before and after coupling of transferrin (Tf) and poly-L-arginine (PR) at pH 7.4. Data are shown as means and standard deviation (n=3). nd: not determined.

| Liposome composition | Physical properties |  |  |
| --- | --- | --- | --- |
| | Particle Size (nm) | PDI | ζ-potential (mV) |
| Plain (empty) | 157.6 ± 1.1 | 0.102 ± 0.013 | 7.16 ± 0.72 |
| Plain (encapsulating pBDNF) | 138.9 ± 0.8 | 0.1065 ± 0.013 | 5.915 ± 0.71 |
| Plain (complexated with pBDNF) | 178.4 ± 0.8 | 0.082 ± 0.033 | -4.18 ± 2.94 |
| Tf-liposomes (encapsulating pBDNF) | 125.1 ± 0.6 | 0.152 ± 0.014 | -8.73 ± 1.13 |
| PR-liposomes (encapsulating pBDNF) | 124.7 ± 0.5 | 0.106 ± 0.014 | 7.83 ± 0.68 |
| Tf-PR-liposomes (empty) | 177.24 ± 75.5 | nd | nd |

As expressed in table 1, the different liposome preparations present diverse physical properties. As expected, the results showed that the incorporation of Tf decreased the zeta potential to a negative value. Anabousi et al. (2005) have demonstrated that using DSPE-PEG_2000_-COOH as linker lipid led to the highest amount of bound Tf, and the negative charge of the ζ-potential indicated that Tf was attached to the liposome surface. On the other hand, usually, the coupling of PR to the liposome surface yields a higher positive charge. Sharma et al. (2012) demonstrated that PR-liposomes presented a ζ-potential of 20.25 ± 3.6 mV, which were stable after 30 days storage presenting ζ-potential of 21.30 ± 3.5 mV (Sharma et al., 2013). Kim et al. (2010) have also shown that PR-PEGylated liposomes displayed ζ-potential of 32.2 ± 3.7 mV. Therefore, considering that the ζ-potential of our PR-liposomes was very similar to that of the plain liposomes, we can suggest that PR-PEG coupling efficiency was either low or did not occur.

Furthermore, we also noticed that the different ways of loading the liposomes with pDNA generated different phisical characteristics in the plain liposomes. The addition of pDNA during liposome formation resulted in posively charged liposomes (Parker et al., 2003). Lipoplexes formulations, in which pDNA was added to the solution containing the already formed liposome (Kim et al., 2010), instead showed a slight negative ζ-potential, suggesting the presence of a higher amount of pDNA on the surface of liposomes. Sakurai et al., (2000) showed that cationic liposomes can aquire a negative charge depending on the amount of plasmid added to the formulation. Thus, it is likely that the pDNA post-added to the liposome formulation modulated the ζ-potential conferring negative surface charge to the plain liposomes.

### Plasmid encapsulation efficiency

The pDNA loading efficiencies for plain, Tf, and PR-liposomes were 44.1%, 27.6%, and 57.5%, respectively. Our results are in agreement with those of Sharma et al. (2012), in which plain liposomes presented an efficiency loading of 35±4.3%, and Tf- and PR-liposomes, encapsulation efficiencies of 33±5.2% and 40±4.1%. However, we observed that several factors interfered with the absorbance measurements. For example, we initially used 1% Triton-x in order to break the liposomes and measure the freed pDNA concentration. However, we observed that the Triton-x interfered with the DNA measurement showing absorbance at 260nm. We replace the Triton-x by methanol (Podesta and Kostarelos, 2009; Zhang et al., 2010), and then, we were able to achieve more reliable results. Still, for further studies, we would like to conduct agarose gel electrophoresis to better access the pDNA content of the liposomes (Kim et al., 2010).

### Gene expression

Plain liposomes and PR-liposomes encapsulating pGFP were used to access the transfection efficiency in neuroblastoma cells (SH-SY5Y) using a live-observing fluorescent microscopy. **Figure 4** shows the bright field representation of the images taken during the experiment. We were not able to observe any fluorescence emission 0, 24, 48 or even 72 hours after adding the liposomes to the cells, indicating that these liposomes did not transfect the cells. The neuroblastoma cells line used in this study are often used to understand neuronal signaling. These cells are typically transfected via the calcium phosphate method or electroporation (Carrì et al., 1997; Hasegawa et al., 2004), but also different reports demonstrated successful transfection with liposomes, indicating that these cells should be sensitive to liposomal transfection (Betz et al., 2003; Obata et al., 2010). Therefore, other factors might have contributed to the failure of the transfection. It is likely, for example, that the cells did not uptake the liposomes or that the purification of the liposomes was not sufficient leading to cytotoxic effect over the cells. In conclusion, further improvement of the liposome synthesis will be necessary to achieve optimal transfection.

**Figure 4.**
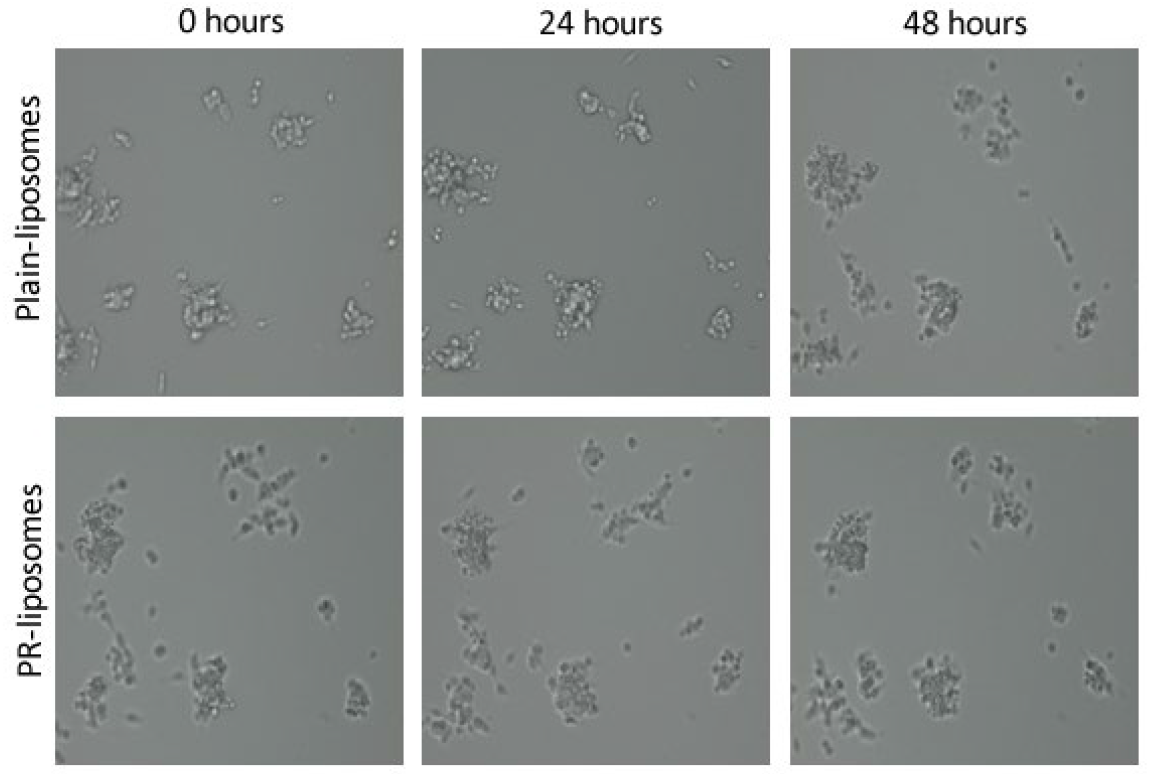
Live-cell imaging of *SH-SY5Y cells transfected by plain and PR-liposomes* using Invitrogen EVOS M7000 Imaging System. Bright-field and fluorescence pictures were taken hourly by the automated microscope. Selected bright-field images at 0, 24, and 48 hours were chosen to represent the study, but fluorescence images did not display any emission.

## Conclusion and perspectives

In this study, we described the preparation of liposomal carriers to promote gene delivery through the blood-brain-barrier. We synthesized plain liposomes containing cationic lipids and PEGylated surface, PR-liposomes containing the cell-penetrating peptide arginine, and Tf-liposomes containing transferrin which promotes receptor-mediated endocytosis. Moreover, we used both conjugates to develop the bifunctional dual-targeted Tf-PR-liposomes. This pilot study revealed the challenges of tagging the liposomes with the ligands Tf and PR. Notably, we encountered several limitations when verifying the coupling efficiency between the DSPE-PEG and the conjugates PR and Tf. Purification of the free peptides from the final liposomal product is a critical step to effectively validate liposome formulations for their clinical use to ensure that the observed therapeutic effects are due to the liposome conjugate and not from non-conjugated contaminants (Manjappa et al., 2011; Nobs et al., 2004). This step in the liposome production will therefore need further analysis.

Once the pBDNF-loaded liposome carrier is well characterized, subsequent studies should evaluate the delivery of specific BDNF gene transcripts to target desired brain regions (Aid et al., 2007; Baj et al., 2011; Pruunsild et al., 2007). Besides vector modifications, gene delivery efficiency can also be dramatically enhanced through strategic modification of DNA composition and conformation, thereby improving bioavailability, biocompatibility, durability, and safety (Foldvari et al., 2016). The use of cell-type-specific promoters will confine the transgene expression to a specific cell type. For example, decreased BDNF levels in the hippocampus characterize the pathology of depression (Castrén et al., 2007; Castrén and Rantamäki, 2010; Dwivedi et al., 2003; Ray et al., 2011). Identification of the subcellular localization of the hippocampal BDNF decrease will help in the desing of a plasmid vector that contains a promoter enhancing BDNF gene expression in that specific cellular subpopulation. In principle, specifically targeting a population of neurons or glial cells might allow to achieve the therapeutic goal without off-target effects (Ingusci et al., 2019). For example, the phosphate-activated glutaminase (PAG) or the vesicular glutamate transporter (vGLUT) promoter ensures ^~^90% glutamatergic neuron-specific expression, whereas the glutamic acid decarboxylase (GAD) promoter ensures ^~^90% GABAergic neuron-specific expression (Rasmussen et al. (2007). Furthermore, the specific cellular location of the astrocyte-specific glial fibrillary acidic protein (GFAP) in the CNS has encouraged its extensive use to target transgene expression to cells of glial origin (Lee et al., 2008). Astrocytes as a target is desirable because this cell is in immediate contact with the BBB; additionally, a study designed by Quesseveur et al. (2013) indicated that BDNF released from astrocytes acts on post-synaptic cells in the hippocampus to stimulate neurogenesis, and mediate related anxiolytic-and antidepressant-like activities.

Combining several physico-chemical and genetic modifications into multicomponent non-viral vectors, such as liposomes, is an ongoing challenge but should prove possible. Optimizing both vector and genetic load provides a powerful tactic for development of effective non-viral gene therapy systems that might lead to a new therapeutic approach to deliver BDNF to the brain. Although further research is necessary to determine the safety of gene therapy and to improve drug targeting, liposome-mediated gene transfer is a promising avenue of research into the treatment of central nervous system diseases.

## Acknowledgement

We would like to thank professor Paola Minghetti (department of Pharmaceutical Sciences from the University of Milan) and professor Gerard Martens (department of Molecular Animal Physiology, Radboud University Medical Center) for providing the expertise and facilities for the experiments described in this study. We would like to also thank Dr. Joanes Grandjean for helpful discussions, suggestions, and critical reading of the manuscript.

